# Progressive *Mycobacterium abscessus* lung infection in C3HeB/FeJ mice associated with corticosteroid administration

**DOI:** 10.1101/418491

**Authors:** Emily C. Maggioncalda, Elizabeth Story-Roller, Nicole C. Ammerman, Eric L. Nuermberger, Gyanu Lamichhane

**Affiliations:** Division of Infectious Diseases, Department of Medicine, School of Medicine, Johns Hopkins University, Baltimore, MD 21287.

**Keywords:** *Mycobacterium abscessus*, pulmonary infection, animal model

## Abstract

*Mycobacterium abscessus* (*Mab*) is a rapidly-growing nontuberculous mycobacterium that is a growing health concern among both immunocompetent and immunocompromised patient populations. It most commonly causes skin and soft tissue or pulmonary infection. As an emerging health issue there is much that still needs to be understood about the infection and its progression to disease. In the context of pulmonary infection, an *in vivo* system of *Mab* infection that permits investigation of host-microbe interactions that result in *Mab* infection and the transition to pathogenesis, and also the evaluation of treatments, is an essential tool that is currently lacking. Here, we describe a system of pulmonary *Mab* infection in the C3HeB/FeJ mouse strain under corticosteroid immunosuppressive therapy that progresses to pathology.

## INTRODUCTION

*Mycobacterium abscessus* (*Mab*) is a rapidly-growing nontuberculous mycobacterium. This environmental mycobacterium, while often an innocuous colonizer, can cause pathology in the lungs, skin, and other soft tissues. In patients with underlying immunodeficiency and/or lung conditions such as cystic fibrosis or bronchiectasis, *Mab* can cause a chronic lung infection^1,2^. Despite the emerging significance of *Mab* pulmonary infections, a mammalian model that recapitulates the process of aerosol infection and subsequent development of pathology in human disease^3–5^ has not yet been developed. Such a model would be an essential tool to study pathogenesis of *Mab* disease and to assess efficacy of treatment regimens.

In recent years, several attempts have been made to develop models of *Mab* infection. Models using hosts such as *Drosophila*^6,7^, zebrafish^8–13^, and *Galleria mellonella* larvae^14^ have been described, but these models do not capture key aspects of pulmonary infection. As mouse models have been used extensively for the study of pulmonary *M. tuberculosis* infection and disease^15–18^, it is relevant to assess if the mouse can serve as a host to develop a model of pulmonary *Mab* disease that is representative of aspects of human infection and pathology. We therefore turned to the mouse to develop a system of pulmonary *Mab* infection that utilizes the aerosol route for infection, and leads to an increase in lung bacterial burden after implantation resulting in the formation of gross lung lesions. Several studies have generated significant insights into infection dynamics of *Mab* in various genetic backgrounds of mice and demonstrated that immunocompetent mice gradually clear the infection, while certain immunodeficient mice can sustain chronic infection. The intravenous route of infection used in four of these studies^19–22^ represents a septicemia model and therefore is unlikely to mimic early stages of host-pathogen interactions that occur during infection via the aerosol route. Similarly, inoculation of *Mab* bolus intratracheally^23–25^ does not adequately mimic the process of aerosol infection. This approach is also highly invasive, requiring survival surgery for each mouse. Using intratracheal infection, Byrd *et al*. reported that *Mab* sustains chronic infection in the lungs of SCID mice but is cleared rapidly in immunocompetent BALB/c mice^23^, while Choi *et al*. observed similar clearance in immunocompetent C57BL/6 mice^24^. Caverly *et al*. demonstrated a relatively stable lung bacterial burden in C57BL/6 mice using a modified fibrin plug model of intratracheal infection, but clearance of the infection did occur in “cystic fibrosis” S489X Cftr mutation mice^25^

Several papers have attempted to bypass these more invasive modes of infection in favor of the aerosolization method, with varying degrees of success. Ordway *et al*. observed that while immunocompetent guinea pigs and mice gradually cleared *Mab* infection, in immunocompromised IFN-γ knockout mice *Mab* established infection in the lungs and spleen and proliferated in these organs^26^. De Groote *et al*. demonstrated in the GM-CSF knockout mouse a relatively stable lung bacterial burden with dissemination to the spleen as well, but the development of lung pathology is likely impacted by the lack of this important immunological differentiation factor^27^. N’Goma *et al*. also utilized the aerosol route of infection to compare the virulence of *Mab* cocultured with amoebae in Balb/c mice, although they did not demonstrate any increase in lung bacterial burden over time in this immunocompetent strain^28^. The same lack of increase in lung bacterial burden after aerosol infection was observed by Le Moigne *et al*. in the “cystic fibrosis” ΔF508 mouse^29^.

Based upon these previous studies, we hypothesized that transient pharmacological-induced immune suppression of immunocompetent mice may allow proliferation of *Mab* during acute stage infection followed by a sustained infection that leads to development of lesions. As mentioned, development of pathology is a valuable metric of infection for our system because of its usefulness in future investigations of *Mab* virulence and host pathogen interactions. We therefore chose C3HeB/FeJ mice as the host based on the increasing evidence that, among immunocompetent mice, this strain most closely mimics the pathology of disease in humans from infection of a related mycobacteria, *M. tuberculosis (Mtb)*^15–17,30^.

Although C3HeB/FeJ mice are not inherently immunocompromised, we hypothesized that we could achieve a sustained infection with associated lesion formation in C3HeB/FeJ mice with the right immunosuppressive regimen. This model also allows for exogenous modulation of immune suppression and, therefore, is nimble to experimental needs. This hypothesis of the use of corticosteroids for immune suppression was based on epidemiological studies describing corticosteroid use as a risk factor for development of invasive infections with rapidly-growing mycobacteria^31–33^. Of note in *in vivo* studies, a study evaluating efficacy of antibiotic treatment against *M. ulcerans*, another nontuberculous mycobacteria, in mice in the setting of concurrent dexamethasone use found that corticosteroid use prior to infection resulted in increased bacterial burdens and higher degrees of pathology compared to control^34^. Additionally, studies using the Cornell model of latent tuberculosis^35^ have assessed rates of reactivation of latent tuberculosis in the setting of corticosteroid use^36^. We applied these concepts by administering corticosteroids to the mice to increase their susceptibility to an initial infection. With the combination of immune suppression and use of C3HeB/FeJ mice we hoped to design a sustained *Mab* pulmonary infection model with corresponding pathology development.

## METHODS

### Ethics

Animal procedures used in the following studies were performed in adherence to the Johns Hopkins University Animal Care and Use Committee and to national guidelines.

### Bacterial Strains and Growth Conditions

*Mycobacterium abscessus* strain ATCC 19977 was used in all studies. *Mab* was cultured in Middlebrook 7H9 broth (Becton&Dickinson 271310) supplemented with 10% ADS (Albumin, Dextrose, NaCl) and 0.05% Tween-80^37^ with constant shaking at 220 RPM in an orbital shaker at 37°C. Organ homogenates were cultured on Middlebrook 7H11 selective agar (Becton&Dickinson 283810) supplemented with 10% ADS, 50 µg/mL cycloheximide (Sigma-Aldrich C7698), 25 µg/mL polymyxin B (Sigma-Aldrich P1004), 50 µg/mL carbenicillin (Sigma-Aldrich C1389) and 20 ug/mL trimethoprim (Sigma-Aldrich T7883). Quantitative and colony forming unit (CFU) counts were determined as previously described^17^, but with enumeration of plates after 5 days of incubation at 37°C.

### Mice, infection and corticosteroid administration

Female Nu/Nu (Nude) mice, 4-5 weeks old, were procured from Charles River Laboratories. Female C3HeB/FeJ mice, 4-5 weeks old, were procured from Jackson Laboratories. All infections were performed by aerosolizing *Mab* culture using a Glas-Col Inhalation Exposure System, as per the manufacturer’s instructions. The infection cycle comprised of 15 minute of pre-heat, 30 minutes of nebulization, 30 minutes of cloud decay and 15 minutes surface decontamination. Corticosteroid regimens and study design for Study 1 and Study 2 are outlined in Tables 1 and 2, respectively. Cortisone (Sigma-Aldrich C2755) and dexamethasone (Sigma-Aldrich D1756) were dissolved in sterile 1x phosphate buffered saline, pH 7.4 (Quality Biological, 114-058-101) and administered by sub-cutaneous injection, 200 µL bolus using 26 gauge syringe, as described previously^34,36,38,39^.

### Study 1

We used Nude and C3HeB/FeJ mice. C3HeB/FeJ mice were divided into two groups. The first group did not receive any corticosteroid. The second group received cortisone daily (7 days/week, 40 mg/kg) beginning two days prior to infection. A cortisone dose of 40 mg/kg and daily dosing frequency were selected based on published usage in *Mtb* infection models^36,38^. The athymic Nude mouse group acted as a comparison to observations reported in the literature^19,20^. All mice were infected simultaneously by aerosolizing 10 mL of *Mab* suspension,A_600nm_= 1.0. Lungs were dissected at twenty-four hours post-infection to enumerate the CFU implanted during infection and then at two, four and six weeks post-infection to determine dynamics of infection. A schematic of this study is outlined in **Table 1**.

**Table 1:**
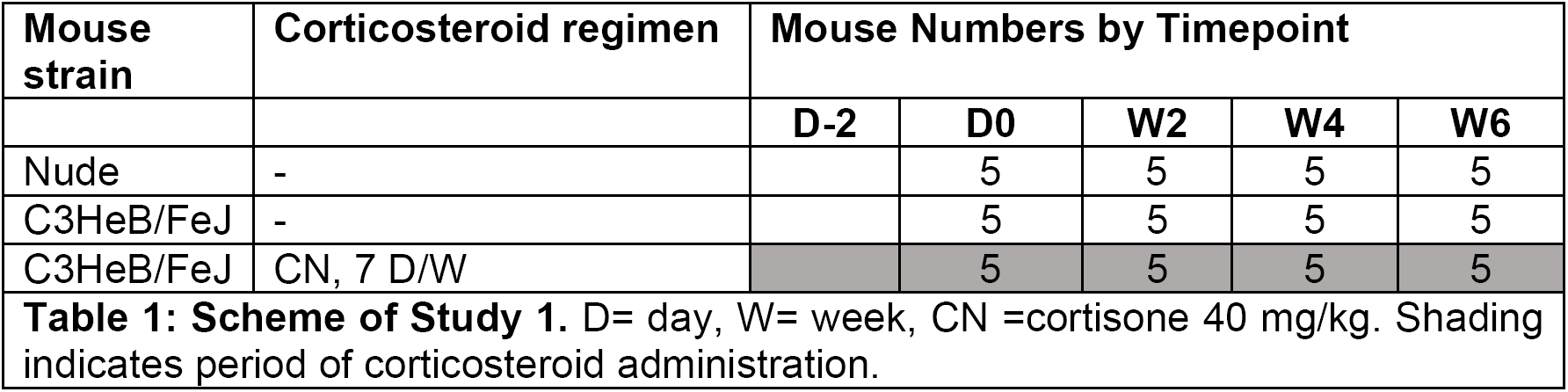
**Scheme of Study 1.** D= day, W= week, CN =cortisone 40 mg/kg. Shading indicates period of corticosteroid administration.

### Study 2

C3HeB/FeJ mice were infected with *Mab* with either a ‘low’ or ‘high’ dose. For low dose, 10 mL of *Mab* suspension at A_600nm_of 0.1 was used. For high dose, 10 mL of *Mab* suspension at A_600nm_of 1.0 was used. In both infection groups there were three corticosteroid regimen groups: no corticosteroid, cortisone 40 mg/kg, or dexamethasone 10 mg/kg. We included dexamethasone with the aim to identify the optimal agent for inducing immune suppression. Cortisone was administered daily beginning two weeks prior to infection, and ending two weeks post-infection. Dexamethasone was administered on Mondays, Wednesdays, and Fridays, beginning two weeks prior to infection, and ending two weeks post-infection. A single dose of cortisone or dexamethasone was administered three days post treatment cessation to taper off the steroid exposure in the mice. Mice were sacrificed twenty-four hours post infection to determine bacterial burden at implantation and then two and four weeks post-infection. CFU determinations in the liver and spleen were undertaken to assess the extent of dissemination and growth in these organs. A schematic of this study is outlined in **Table 2**.

**Table 2:**
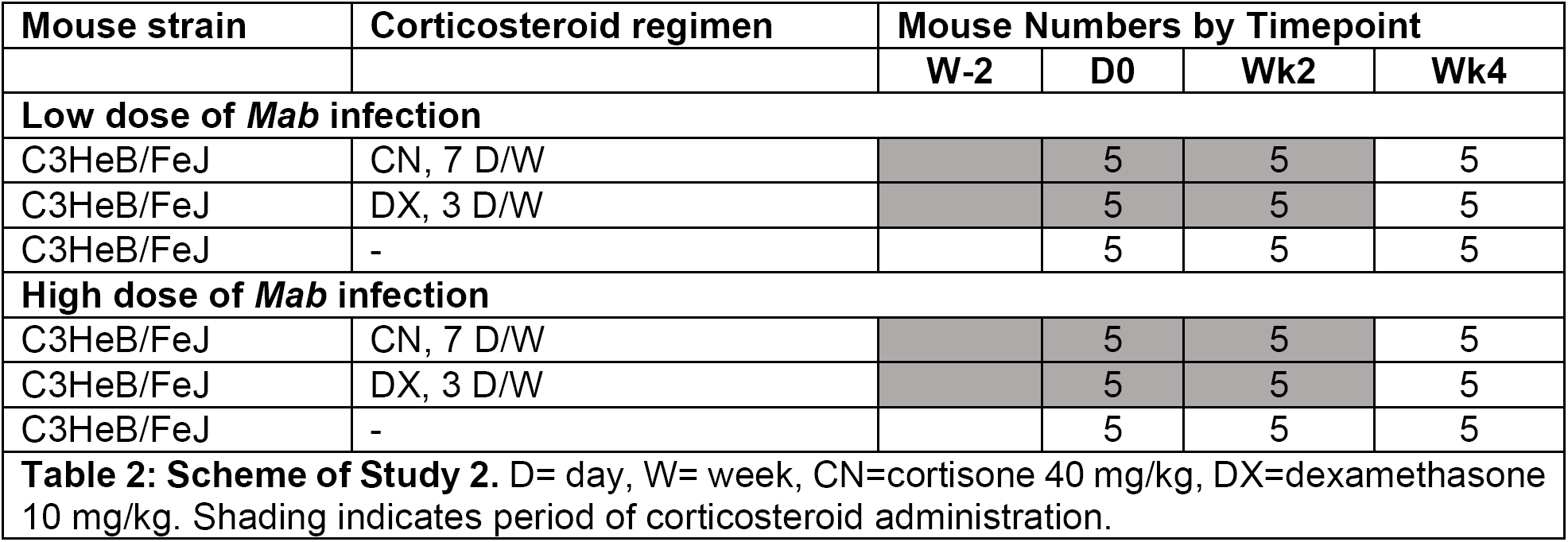
**Scheme of Study 2.** D= day, W= week, CN=cortisone 40 mg/kg, DX=dexamethasone 10 mg/kg. Shading indicates period of corticosteroid administration.

## RESULTS

### Study 1

The burden of *Mab* in the lungs of mice are shown in **Figure 1A**. On the day after infection, similar *Mab* CFU counts were found in the lungs of the three groups of mice: Nude, untreated C3HeB/FeJ and C3HeB/FeJ treated with cortisone for two days. In the lungs of untreated C3HeB/FeJ mice, *Mab* burden declined gradually initially (mean CFU of 3.4 log_10_ to 2.8 log_10_ from day zero to week two) and then more rapidly after two weeks (mean CFU of 2.8 log_10_ to 0.3 log_10_ from week two to week four). Similarly, *Mab* burden also declined in the lungs of Nude mice, but at a more gradual rate. The mean CFU from day zero was 3.6 log_10_, while at week two it was within one standard deviation at 3.5 log_10_, but then declined to 2.7 log_10_ at week four and 1.1 log_10_ at week six. In C3HeB/FeJ mice receiving cortisone, the lung burden was relatively stable for the first two weeks (mean CFU of 3.6 log_10_ at day zero and 3.9 log_10_ at two weeks), followed by a rapid increase thereafter to 4.8 log_10_ at week four and 6.6 log_10_ at week six. At week six one mouse that received cortisone developed a pulmonary lesion (**Figure 1B**).

**Figure 1:**
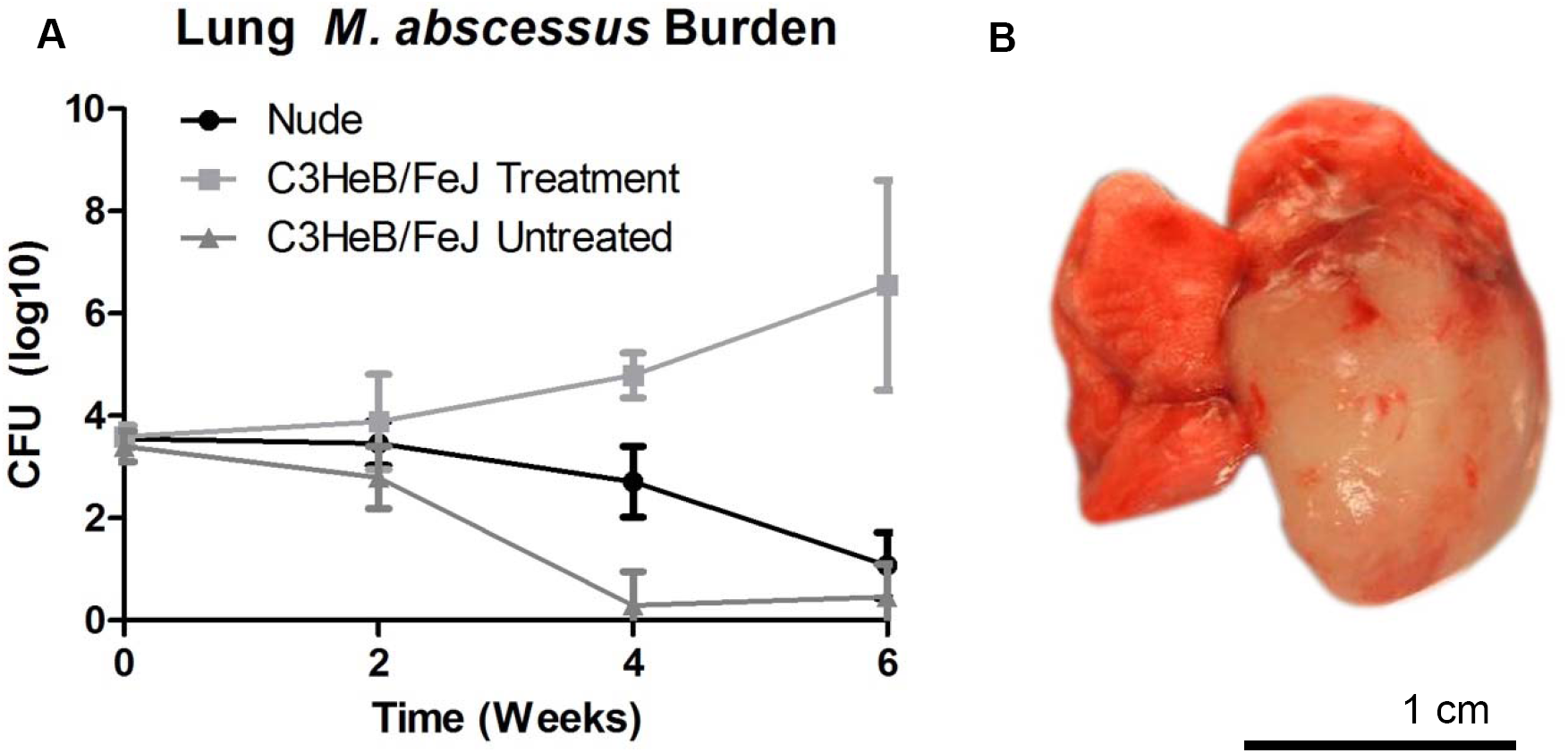
**(A)** Study 1 Lung *Mab* burden, n=5, mean ± SD. **(B)** Lungs of a C3HeB/FeJ mouse treated with cortisone, at 6 weeks post infection.

### Study 2

We initiated a new study with the aim to reproduce and verify *Mab* proliferation and lesion formation observed in the lungs of C3HeB/FeJ mice in Study 1. Compared to Study 1, we administered immunosuppressive treatment for a longer period prior to infection with the intention to decrease the variability in CFU burden observed at two weeks, which was thought to be due to incomplete immunosuppression at infection and in the weeks following.

*Mab* burden in the lungs are shown in **Figure 2**. These infections resulted in an initial pulmonary implantation of 3.9±0.11 log_10_ and 4.4±0.10 log_10_ CFU for low and high *Mab* doses, respectively, averaged across all groups. Two weeks after infection, we observed a rapid increase in bacterial burden in the lungs of mice treated with cortisone or dexamethasone. For cortisone regimen mice with low dose and high dose infection, this meant an increase to 6.8 log_10_ and 6.7 log_10_ mean CFU, and for dexamethasone regimen mice with low dose and high dose infection an increase to 8.7 log_10_ and 7.9 log_10_ CFU. In the untreated mice at this same timepoint we observed a decrease to a mean CFU of 2.8 log_10_ in the low dose infection and 4.1 log_10_ in the high dose infection. We also observed bacterial dissemination to the liver and spleen in some of the mice (**Tables 3, 4, and Supplement Table 1**).

**Figure 2:**
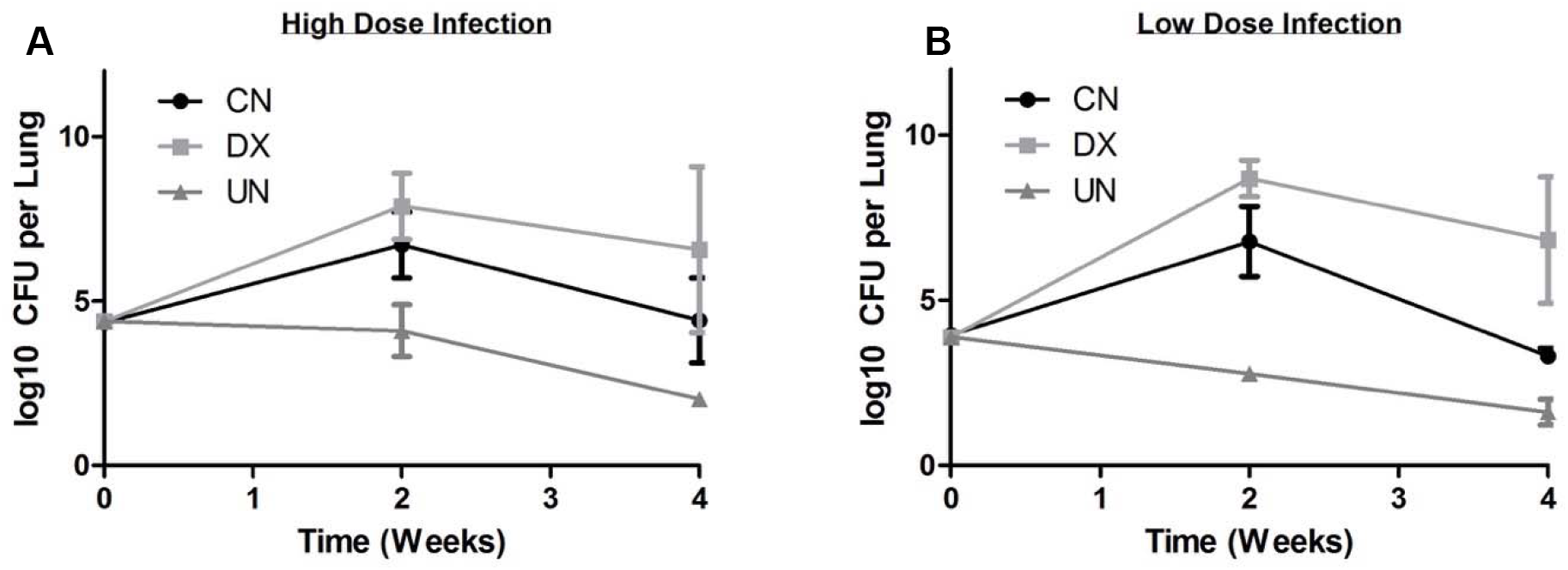
Study 2 *Mab* lung burdens. **(A)** Burden in the lungs of mice infected with a high dose of *Mab*. **(B)** Burden in the lungs of mice infected with a low dose of *Mab*. For all timepoints, n=5 and mean ± SD are reported. CN= cortisone regimen, DX= dexamethasone regimen, UN= untreated.

Shortly after discontinuing treatment, three mice infected with a high dose of *Mab* and treated with cortisone died. Similarly, in groups infected with high and low doses of *Mab* and treated with dexamethasone, three and nine mice died, respectively. There were no deaths in mice infected with low dose *Mab* and treated with cortisone.

At the week four time point, we observed a decrease in the average *Mab* burden in the lungs of mice in all groups. In untreated mice, lung bacterial burden decreased further to a mean CFU of 1.6 log_10_ CFU for low dose infection and to a mean CFU of 2.0 log_10_ for high dose infection over this two week period. For cortisone regimen mice with low dose and high dose infection mean CFU decreased to 3.3 log_10_ and 4.4 log_10_, and for dexamethasone regimen mice with low dose and high dose infection they decreased to 6.8 log_10_ and 6.6 log_10_ mean CFU. The bacterial burden in the liver and spleen, however, continued to increase or remained stable from week two (**Tables 3 & 4**).

**Table 3:**
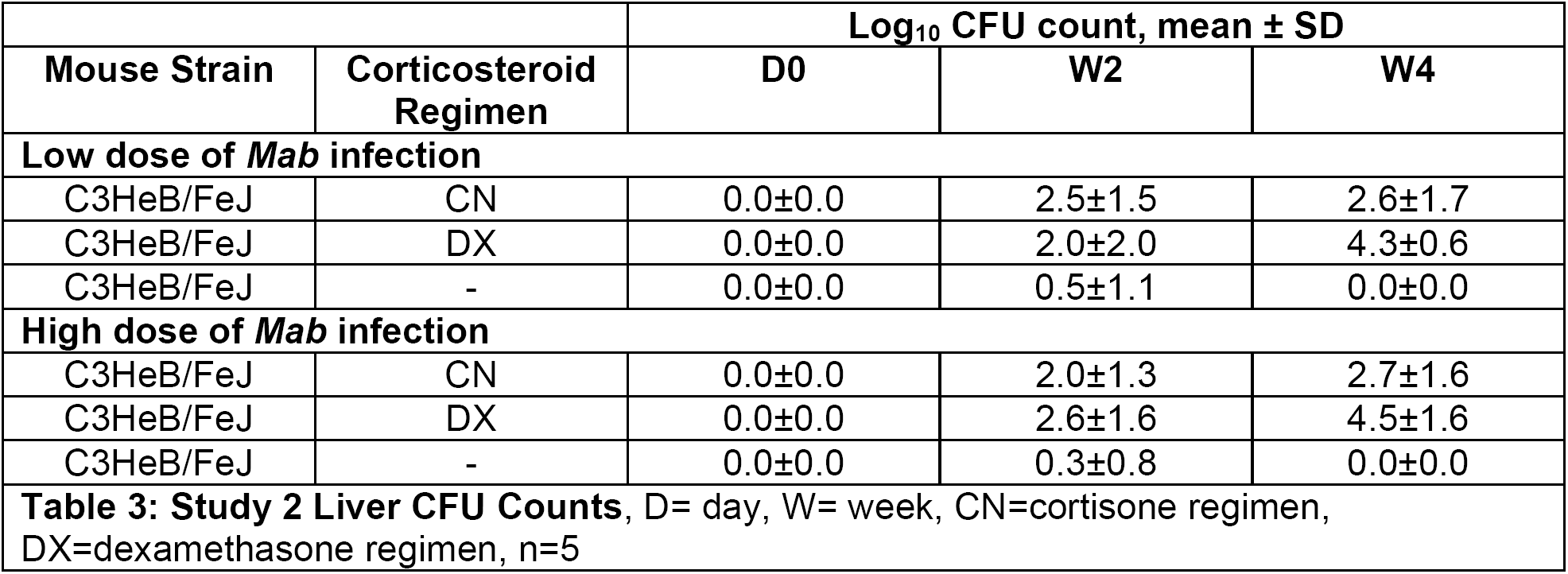
**Study 2 Liver CFU Counts**, D= day, W= week, CN=cortisone regimen, DX=dexamethasone regimen, n=5

**Table 4:**
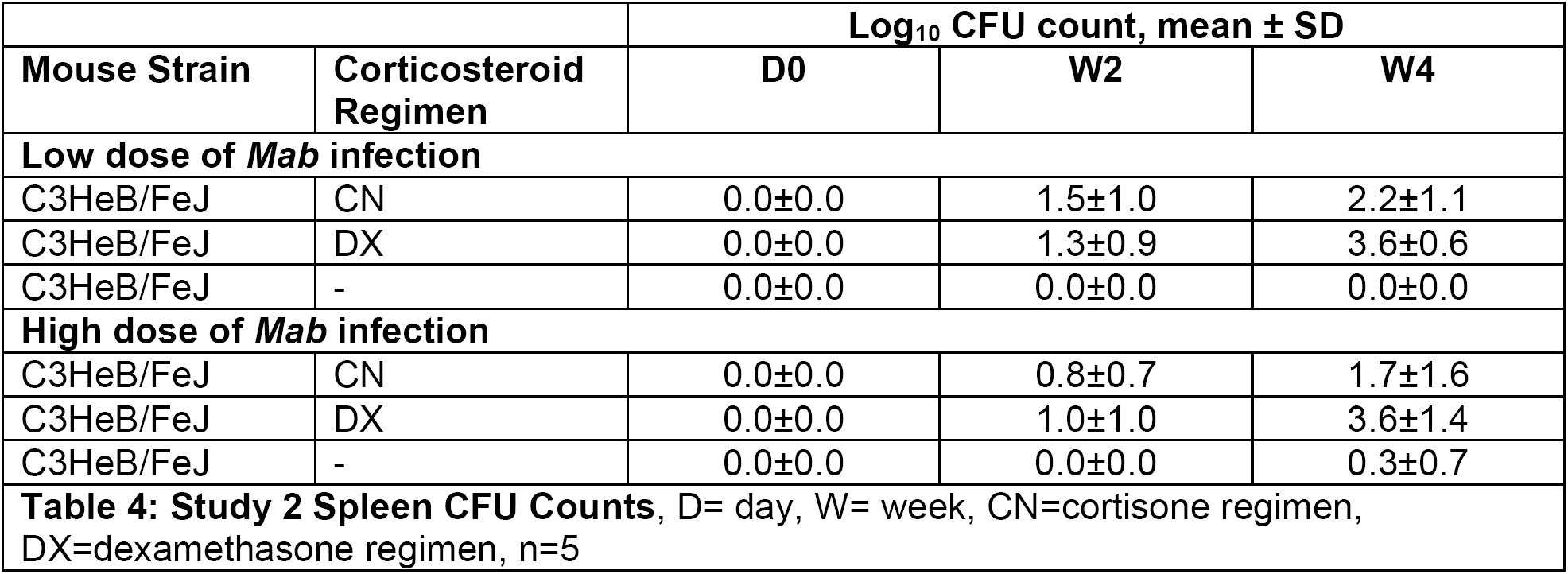
**Study 2 Spleen CFU Counts**, D= day, W= week, CN=cortisone regimen, DX=dexamethasone regimen, n=5

Of note there was an increase in the variability of the lung bacterial burden at this timepoint compared to earlier timepoints, specifically in the dexamethasone regimen groups with both low and high infection doses. Data detailing the weights of all mice in this experiment along with their respective bacterial burdens at the sampled sites (**Supplement Table 1**) indicates that there appears to be a split in CFU burden that corresponds inversely with the weight of the animal, i.e. lower weight mice had higher bacterial burdens. In particular the dexamethasone regimen group at the week four timepoint mice with lower weights had higher lung CFU burdens and vice versa, resulting in the observed variability.

Observations of gross pathology of the lungs of mice at week 4 are shown in Figure 3. In the mice infected with low and high doses of *Mab* but not treated, the gross appearance of their lungs resembles the lungs of uninfected age-matched mice. In the cortisone treatment groups, similar levels of pathology, in terms of size, number and appearances of lesions in the lungs, were observed in both high and low dose infection groups. The lesions in the lungs of dexamethasone treated mice were larger in size and more numerous. The main factor that differentiated the levels of gross pathology observed in the lungs of mice in high or low infection dose groups was that the lungs of mice in the high dose infection group developed such a degree of inflammation and necrosis that they had fused to the thoracic wall, which made their intact extraction difficult. The levels of lung pathology observed qualitatively appear to correspond with the levels of bacterial burden in these groups over the course of the infection.

**Figure 3:**
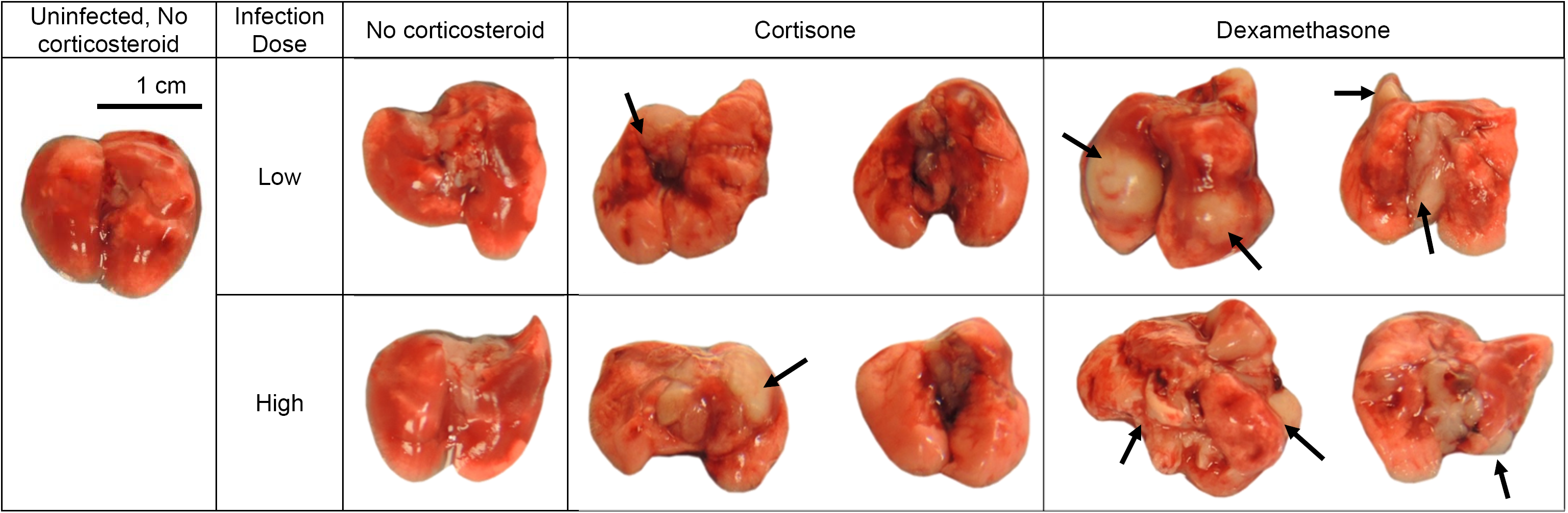
Gross pathology of lungs of mice at week four timepoint, Study 2. Lungs are representative of their group. Arrows indicate observable lesions.

## DISCUSSION

Our aim was to develop a system of pulmonary *Mab* infection in mice that utilizes the aerosol route of infection, leads to an increase in lung bacterial burden after implantation, and develops pulmonary lesions due to infection. The studies of *Mab* infection that have been performed in mice thus far have unfortunately failed to achieve these goals in an immunocompetent mouse strain. The study presented here describes a sustained *Mab* pulmonary infection via aerosolization in an immunocompetent mouse strain with corticosteroid-induced immune suppression.

Our first study affirmed that cortisone treatment in mice allows for a gradual increase in bacterial burden (**Figure 1A**). However, there was large variability in the bacterial burden at two weeks post-infection and a more uniformly higher bacterial burden was not observed until the week four timepoint. The variability in bacterial burden suggested that a longer duration of corticosteroid treatment prior to infection may be beneficial in increasing the initial bacterial burden in the lungs. Alternatively, this variability may also be explained by the mouse strain itself. C3HeB/FeJ mice are known to have variable infection outcomes when infected with *M. tuberculosis*^16,17^. To test this first hypothesis, we extended the corticosteroid regimen prior to infection in our second study. Here we observed the CFU in the lungs of mice treated with either cortisone or dexamethasone rapidly increased by the week two timepoint and did demonstrate decreased variability at this timepoint. We also observed dissemination to the liver and spleen in corticosteroid regimen mice, which did not occur to the same extent in untreated mice. As noted previously, several of the corticosteroid regimen mice subsequently died following cessation of steroids. However, these mice subjectively appeared to be sicker, of lower body weight, and seemed likely to die from the infection even prior to stopping treatment suggesting that the level of immune suppression was perhaps too high, especially in the dexamethasone-treated group.

Following cessation of corticosteroid administration, mice may have experienced some level of immune reconstitution, as evidenced by the decreased bacterial burden in the lungs observed across all groups at the week four timepoint. The rate of clearance observed after stopping corticosteroid administration appears to be gradual enough that experiments investigating virulence of *Mab* or treatment regimens against *Mab* would be able to be implemented with this system to examine changes in bacterial burden over time. Interestingly, despite the decrease in lung CFU, the bacterial burden in the liver and spleen remained stable or continued to increase. Dissemination of *Mycobacterium chelonae,* another rapidly growing mycobacteria, to skin and soft tissue has been associated with chronic use of low dose corticosteroid^31^. Additional studies to optimize the immunosuppressive regimen and initial infectious dose are currently underway in order to maximize the period of time the mice maintain a high bacterial burden, while minimizing the mortality.

Although further studies are needed to validate an aerosolized mouse model of a sustained *Mab* pulmonary infection, these preliminary infection studies are quite promising, and may be used as a guide for other investigators working on the development of mouse models of *Mab* infection and disease.

## ACKNOWLEDGMENT

This work was supported by the Cystic Fibrosis Foundation grant. ESR was supported by the NIH T32 AI007291. The content is solely the responsibility of the authors and does not necessarily represent the official views of the Cystic Fibrosis Foundation or the National Institutes of Health.

